# Secundary Structure of Physicochemical Clustered Proteins

**DOI:** 10.1101/2023.05.30.542924

**Authors:** Igor Nelson

**Affiliations:** MsC. Computational Biology, University of Coimbra, Coimbra, Portugal

**Keywords:** Bioinformatics, Secondary Structure, Protein Structure Prediction, Deep Learning, Convolutional Neural network (CNN), Machine learning

## Abstract

Diverse methods have been proposed for protein secondary structure prediction. However, such task still presents a challenge in bioinformatics. In this article various of these methods are implemented and analysed. First, a baseline using Support Vector Machine. Then a convolutional neural network (CNN), a Long Short-Term Memory (LSTM) and a strategy of Ensembling both of these methods. Lastly, a novel technique Secundary Structure of Physicochemical Clustered Proteins (SSPCP) is proposed, which combines multiple CNNs trained accordingly to a protein feature clustering and combined using a neural network. The rationale behind SSPCP is that amino acids from proteins which have similar physicochemical characteristics should have the same secondary structure prediction for similar amino acids, but amino acids from differing proteins might have different structures. All of these methods use as features PSSM matrices extracted from PSIBLAST. For performance evaluation, 25pdb dataset was split into training and validation and the same subsets were used on all these methods achieving the Q3 score of CNN: 70.11%, LSTM: 69.25%, Ensemble: 70.71%, SSPCP: 70.91%.

The experimental results show that the features extracted from clustering of physicochemical properties of proteins seem to improve the accuracy of highly specific CNN models for accurate protein secondary structure prediction.

## I. Introduction

Proteins are the basis for life playing a key role in almost every biological process and providing accurate cellular and molecular functioning. Their functions are determined by specific spatial structure and they can be composed by 20 different amino acids. These composing amino acids are connected in sequence with each carboxyl group of one amino acid forming a peptide bond with the amino group of the next one.

The biological function of proteins is highly related with their tertiary structure, which is in turn believed to be determined by its amino acid sequence via the process of protein folding^1^. In this way, the correct prediction of protein structure is crucial for understanding protein functions and propose new drugs. However, experimental methods such as X-ray or Nuclear Magnetic Resonance cannot meet the actual demand, and thus accurate computational methods are required.

These computational methods have shown interesting results. However, they still present a challenging task in computational biology and proteomics. Actually, predicting structures from protein sequences is one of the most challenging tasks in computational biology^2^. It is hard to predict protein structure directly from amino acid sequence and so the intermediate step of secondary structure prediction has been studied. Other than to predict protein structural classes, secondary structure can be also used to study protein function, regulation and protein interactions.

Secondary structure refers to the local conformation of protein’s polypeptide backbone. The Dictionary of Secondary Structure of Proteins (DSSP)^3,4^ was designed by Wolfgang Kabsch and Chris Sander to standardize secondary structure assignment. DSSP is a database of secondary structure assignments for all protein entries in the Protein Data Bank (PDB)^5^. DSSP is also the program that calculates DSSP entries from PDB entries yet not predicting secondary structure. DSSP automatically assigns secondary structure into eight states (H, E, B, T, S, L, G, and I) according to hydrogen-bonding patterns. These eight states are often further simplified into three states: α-helix, β-sheet and coil.

The metric most used to evaluate secondary structure classifiers is the Q3 accuracy which is no more than the accuracy for a three classes problem measuring the percentage of residues correctly predicted for each class over the overall number of predictions.

The first work related with secondary structure prediction was published in 1951 when Pauling and Corey predicted helical and sheet conformations for protein polypeptide backbones^2^. Since then, diverse methods have been proposed to predict secondary structure from amino acid sequence^6–11^.

From these works it was possible to see that, in general, statistical and physicochemical features related with the amino acid sequences have a relatively low accuracy rate in comparison with other methods, being usually no higher than 65%^9,11^.

One of the best improvements in this field came from the usage of Position specific scoring matrix (PSSM)^12^ usually calculated with on PSIBLAST^13^ which reflect the information of sequence evolution, amino acid conservation and mutation. By combining these features with machine learning methods, it was possible to improve accuracy to more than 70%^7,10,14–18^.

Up to recent years, methods with best performance consist on using deep learning strategies. These methods usually provide all the information to neural networks and let artificial intelligence build intermediate features as well as deciding which of them to use in order to achieve the best performance. The best-known deep learning results have achieved 82–84% accuracy^2^.

However, is important to note that even for biochemists there are no clear boundaries between helix and coil neither sheet and coil states and thus structural homologies may differ by about 12%. This creates a theoretical upper limit of 88% for the secondary structure classification task.

In this article three methods are implemented regarding secondary structure prediction. In the first one, a baseline Support Vector Machine (SVM) with a Radial Basis Function (RBF) kernel. The second one using the strategy proposed by by Jinyong Cheng et al.^19^ based on the integration of a Convolutional Neural Network (CNN) and Long Short-Term Memory (LSTM). Lastly, our proposed method Secundary Structure of Physicochemical Clustered Proteins (SSPCP), a specific method that combines multiple CNN models trained accordingly to a previous protein feature clustering and a combining neural network. All of these methods use PSSM matrices extracted from PSIBLAST.

## II. Methods

### A. Starting Dataset

The 25pdb dataset was used to train and evaluate the performance of the proposed methods. As the name suggests the homology of protein sequences which compose this dataset if of less than 25%. 25pdb is composed of 1673 proteins. Regarding its structures 443 are all-α, 443 all-β, 346 α/β and 441 α + β.

### B. DSSP

For this study these classes were not used as target output. Instead, secondary structures were extracted from related PDB files using the DSSP software. Note that in this process it was not possible to extract secondary structures for 29 of these proteins. While in the standard 25pdb dataset each sequence corresponds to a unique expected class such as all-α, here each amino acid of the protein was assigned a secondary structure label. The very short summary of the DSSP output label is: H = α-helix; B = residue in isolated β-bridge; E = extended strand, participates in β ladder; G = 3-helix (310 helix); I = 5 helix (π-helix); T = hydrogen bonded turn and S = bend. However, for the sake of structure classification These 8 types of structures can be further converted into 3 types of structures. Here the strategy chosen was to merge G, H and I into H, B and E into E, and the others into C.

As 25pdb dataset presents subparts of proteins and their corresponding structure, some proteins were found repeated, simply because they might have different structures assigned to different parts. Those proteins were only considered once for this study. Also, it was not possible to apply these DSSP procedure to some proteins. Finally, sequences with less than 30 amino acids were removed. The final number of proteins in the dataset was of 1427.

In similar works it is a common practice to use cross-validation in order to minimize the impact between the training set and the test set, however because of time restrains we only performed train test split with 70%/30%. As so, the training part consisted on features extracted from 413.738 amino acids and the testing part consisted in 153.679 amino acids from independent proteins.

### C. PSSM

Using the amino acid sequences present in the PDB files we then run PSIBLAST on all proteins in order to obtain PSSM information. These matrices reflect sequence evolution, amino acid conservation and mutation for present amino acids. Here the method of choice was to use Swissprot as the database and, blossom62 as the evolution matrix with a threshold of 0.001 and 3 iterations. The outputs are a 20*N dimensional matrix for each protein, 20 the possible amino acids and N the number of amino acids in protein sequence.

Then, a sliding window of consecutive amino acids was applied in order to obtain sequence information and predict the secondary structure of the central residue. Each residue was encoded by a vector of dimension 20 × w, where w is the sliding window size. The window was shifted from residue to residue through the protein chain. When the sliding window was at the front end and or back end of the sequence, the window was filled with zeros. The matrices can then be used as input for machine learning and deep learning strategies. These features were converted into a one-dimensional vector for the machine learning methods and in two-dimensional matrices for the CNN models. For a value w=13 that corresponds to a vector of 260 (13*20) features or a 2D array of 13*20 for each amino acid.

### D. Protein Clustering and to Physicochemical Properties

Since amino acids have physicochemical properties, Shen et al.^20^ proposed the categorization of amino acids to reduce the vector space dimensionality. In his work, it was suggested that all amino acids could be binned in seven different categories, as amino acids within the same category would most likely result in synonymous mutations, given their similarities. The substitution table used by Shen et al.^20^ is shown in Table I.

**Table I.**
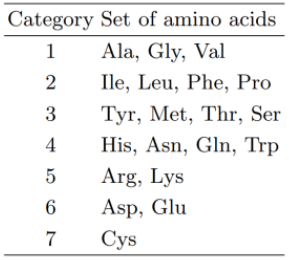
Seven Groups of Amino acids according to Shen et al.^20^

In the present work the same table was applied in *Method 5* in order to extract features and perform protein clustering according to its length and composition. The rationale is that amino acids from proteins which have similar physicochemical characteristics should have the same secondary structure, but amino acids from differing proteins might have different structures. These protein physicochemical characteristics would be related with each amino acid secondary structure.

### E. Implementation Overview

Here, five different strategies were implemented and tested using features extracted with the following methods:

*Method 1 – Baseline Approach:* First analysis of the data and implementation of a simple classifier using SVM and RBF;

*Method 2 – CNN:* Implementation and study of valid CNN model;

*Method 3 – LSTM:* Implementation and study of a valid LSTM model;

*Method 4 –* Ensembling of optimized CNN and LSTM methods into a final classification strategy;

*Method 5 – SSPCP:* In this method proteins are clustering according to physicohemical properties, assigned to different CNN classifiers and then a neural network is used to combine outputs from all these classifiers.

## III. Results And Analysis

### A. Method 1 – Baseline Approach

The first experiments for the baseline approach were made by using a Support Vector Machine (SVM) with a Radial Basis Function (RBF) kernel. SVM classifiers have been used successfully in protein secondary structure prediction^21,22^. The RBF kernel require two parameters: *C* and *gamma*. Intuitively, the *gamma* parameter defines how far the influence of a single training example reaches, with low values meaning *far* and high values meaning *close*. The *gamma* parameters can be seen as the inverse of the radius of influence of samples selected by the model as support vectors. The C parameter trades off correct classification of training examples against maximization of the decision function’s margin. For larger values of C, a smaller margin will be accepted if the decision function is better at classifying all training points correctly^23^.

Since studying the effect of these parameters in the classification task can take a long time, a subset of 60 proteins was created in order to choose adequate parameters for the remaining classification tasks. The best parameters found were then used for the following SVM-RBF classifiers.

In Figure 1 there is a study of the best parameters for *C* and *gamma*. For the given subset the best parameters were *C*: 10.0 and *gamma*: 0.001 and so they were used for the remaining SVM-RBF implementations in this work.

**Figure 1.**
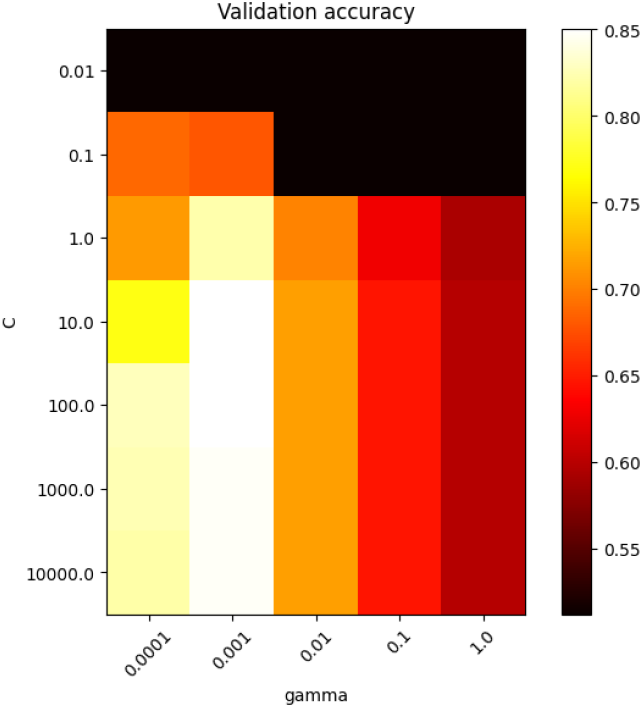
A study for SVM-RBF kernel C and gamma parameters.

Afterwards, using these parameters, it was useful to evaluate the effect of using different values for amino-acid neighbourhood. For that, 3 values (w=11, w=13 and w=15) were applied to a subset of 100 proteins from the 25pdb dataset. Note that these results might not extrapolate to the complete dataset, but due to limited time available for this project it was impossible to evaluate the whole dataset with SVM-RBF.

As reported in previous works^19,24^ w=13 was the best choice for neighbourhood length achieving 0.75 Q3 score in this subset with 100 proteins. Note that this performance would drop if the method was applied to the whole dataset. w=13 was then applied the following implementations in this work.

After that, an investigation was conducted into how similar features extracted with this method would be in relationship to each other. In Figure 3 is possible to see the results of hierarchical clustering in a form of a dendrogram. Most amino acids are grouped together in right hand side of the image.

**Figure 2.**
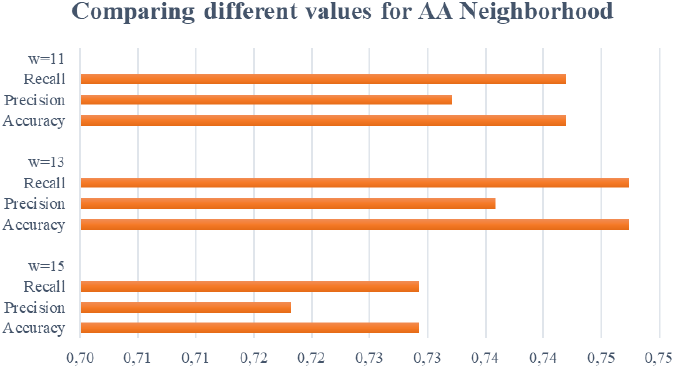
Comparing different window lengths for amino acid neighbourhood.

**Figure 3.**
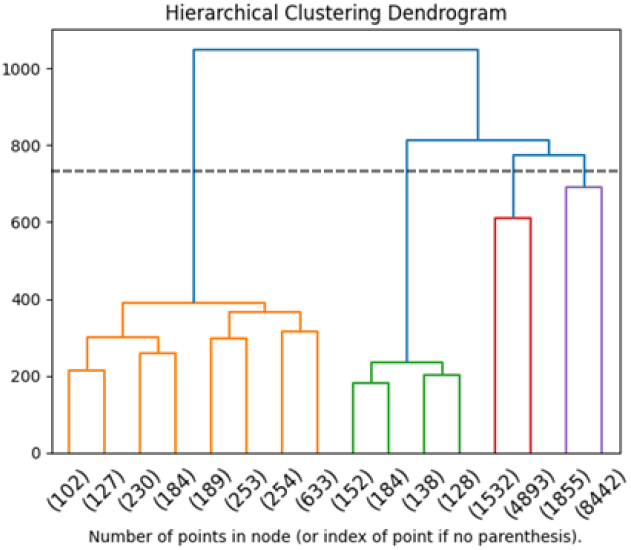
Hierarquical clustering of PSSM features for each amino acid with a window w=13.

As suggested by the traced line in Figure 3, 4 clusters of data were created in order to check how the general classifier trained for 100 proteins was performing for each one this groups of data. In Figure 4 there is a representation on the expected classes for each one of this clusters.

**Figure 4.**
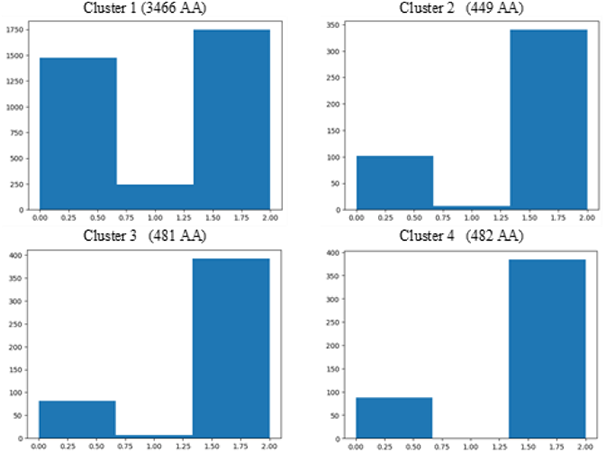
Expected classes per each one of the 4 clusters. 0:coil, 1:strand, 2:helix. AA stands for Amino Acids present in each cluster.

**Figure 5.**
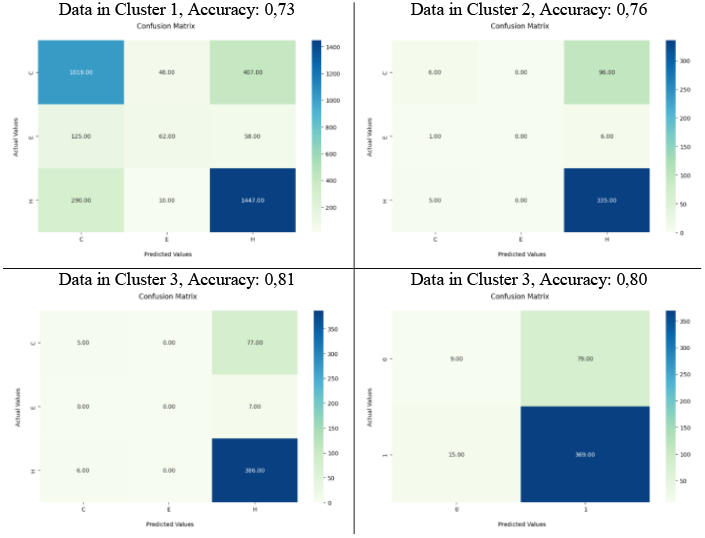
Accuracy of the general classifier in each one of these clusters.

During the implementation there was expectation regarding whether each cluster would be highly correlated with a given type of secondary structure, however that correlation was not found in data. It is important to refer that Cluster 1 shows a different profile when compared with others. It had almost as many expected coils as helices. All clusters had a significantly lower number of strands. Clusters 2, 3 and 4 had overall similar profiles. Here is important to notice that maybe for future works it would be interesting to find datasets with are more balanced in expected classes, since there is a huge discrepancy between expected helix and coil classes and that can limit the classification task.

An evaluation was made into how the general classifier build for 100 proteins would be performing for each one of these clusters). Here it was possible to find that it had a Q3 of 0.73 for the Cluster 1, the one with most proteins; 0.76 for Cluster 2; 0.81 for Cluster 3 and 0,80 for Cluster 4. In Cluster 4 only samples from two classes (coil and helix) were present. It was not possible to find an interesting correlation between the hierarquical clustering of features and the expected output and so this approach wasn’t further explored.

### B. Method 2 – CNN

CNNs have gained attention of scientific community in recent years because they have the ability of being some extent invariant to shift, scale, and distortion. These characteristics make them good methods to problems related with speech^25,26^ and image recognition^27^. These properties also make them a good candidate to protein secondary structure prediction. In recent works, authors used CNN for this problem with great success^28,29^. A CNN implementation was tested in this project following the structure suggested by Jinyong Cheng et al.^19^.

In order to test this structure, in this work 3 models were built and tested changing the number of neurons in the Convolution Layers and in the Dense Layer as shown in Table II. These models were tested on the 100 proteins subset as referenced in above section.

**Table II.** Testing Different Models for CNN.

|  | ConvLayer1 | ConvLayer2 | DenseLayer | Q3 Score (%) |
| --- | --- | --- | --- | --- |
| **Model 1** | 250 | 50 | 25 | 74,57 |
| **Model 2** | 500 | 100 | 50 | 75,78 |
| **Model 3** | 600 | 125 | 60 | 74,26 |

Also kernel size can influence the performance of CNN. 3 models were built changing this parameter for the first Convolution Layer as shown in Table III.

**Table III.** Testing different kernel sizes.

|  | Kernel Size | Q3 Score (%) |
| --- | --- | --- |
| **Model 1** | (3,3) | 74.61 |
| **Model 2** | (5,5) | 75,78 |
| **Model 3** | (7,7) | 69.47 |

From the Q3 score of both tests, the best parameters were obtained from models 2. After testing this models CNN was set as a first convolution layer with 500 convolution kernels of 5*5, and step size of 1 by 1. The next layer was a 2x2 max-pooling layer, which step size is 2*2. The second convolution layer has 100 convolution kernels with the volume of 2*2 and step size of 1*1. Finally, a Fully Connected Layer with 50 neurons and an Output Layer with 3 neurons (using *Softmax activation*). The CNN was trained with Adaptive moment estimation (Adam) optimizer and Categorical Cross-Entropy loss function for 5 epochs.

By using this method, the final classification Q3 on independent testing part of the whole dataset was of **70.11%**. Results from this method were not as good as supposedly found reported in the previous work.

### C. Method 3 – LSTM

LSTM is an extension of the recurrent NN that can deal with remote dependence between sequences and avoid the common problem of gradient disappearance. These networks are usually applied to natural language processing^30^ because they excel at tasks such as extracting meaning from words or finding relationships regarding the order of input. As in text recognition the order of words is of extreme importance for the meaning of phrases so in secondary structure prediction the order of amino acids might be related with protein structure. And this may happen due to remote interactions between amino acids.

LSTM was implemented as following the model described by Jinyong Cheng et al.^19^. Some tests were made with subset of 100 proteins from the dataset in order to choose good parameters for the model. In Table IV the results obtain while changing these parameters are shown.

**Table IV.** Testing different amounts of hidden units in LSTM Model.

|  |  |  |  |  |  |  |
| --- | --- | --- | --- | --- | --- | --- |
| Units in Hidden Layer 1 | 200 | 400 | 600 | 400 | 400 | 400 |
| Units in Hidden Layer 2 | 800 | 800 | 800 | 400 | 600 | 800 |
| Q3 Score (%) | 61,96 | 60,05 | 58,89 | 59,05 | 65,59 | 60,05 |

After testing, the number of hidden units in the first layer was set to 400 and in the second layer was set to 600. The third and the fourth layer are the dense layer composed by 100 neurons and the output layer composed by 3 neurons. Similarly, to CNN, LSTM was trained with *adam* optimizer and Categorical Cross-Entropy loss function for 5 epochs.

The final classification Q3 on independent testing part of the whole dataset was of **69.25%**. Also here, results from this method were not as good as supposedly found reported in the previous work, which might happen because of discrepancies in proteins used to train and test.

### D. Method 4 - Ensembling of CNN and LSTM

The method proposed by Jinyong Cheng et al.^19^ is then composed by a method of ensembling results from these two classifiers (CNN and LSTM) (Figure 6). This strategy tends to perform better when there are different outputs for the predictive results from each individual model. The output probability is given by the probability of the two classifiers by:

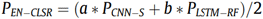

Where a = 0.42 and b = 0.58 which were calculated experimentally by the authors in previous work.

**Figure 6.**
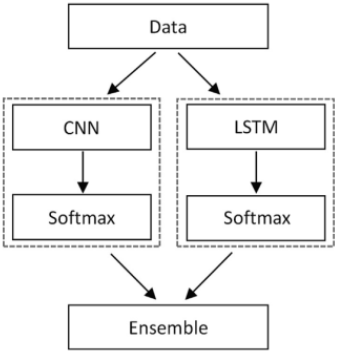
– Ensabling method proposed by Jinyong Cheng et al.^19^

By using this method, the final classification Q3 on independent testing part of the whole dataset was of **70.71%** which suggests combining multiple kinds of classifiers might provide more accurate ways of predicting secondary structure, since there can be multiple reasons why these conformations are present in each amino acid.

### E. Method 5 – SSPCP

The method proposed in this work explores a more complex approach. The first step consists of using the same amino acid substitution table as proposed used by Shen et al.^20^ to categorize amino acids by their physicochemical properties as shown in Figure 7. Then for each protein a feature vector is built combining its length and the counts of each amino acid type from these 7 types, see Table V. These Protein features are then normalized using the standard scaling method.

**Table V.** Features extracted from each protein. a: number of amino-acids. c: percentage of group 1 amino acids regarding whole protein sequence, c: percentage of group 2 amino acids, and so on.

| Features: [a, b, c, d, e, f, g, h] |
| --- |
| Example: [451.00, 0.23, 0.22, 0.21, 0.15, 0.07, 0.11, 0.01] |

**Figure 7.**
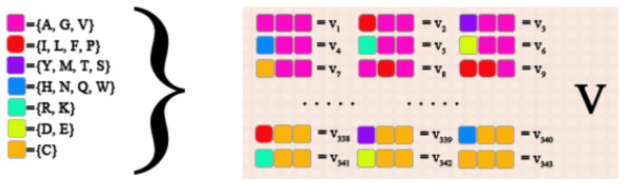
According to Amino acid type features are built in order to characterize each protein. Extracted from Shen et al.^20^ Source PNAS: http://www.pnas.org/content/104/11/4337/suppl/DC1.

After that, a Principal Component Analysis (PCA) transformation is then applied to these features using 2 components. These components are then clustered together. In Figure 8 you can see a plot representing in different colours proteins assigned to each cluster. It is expected that within these clusters proteins are similar regarding protein physicochemical properties and that it might influence secondary structure prediction.

**Figure 8.**
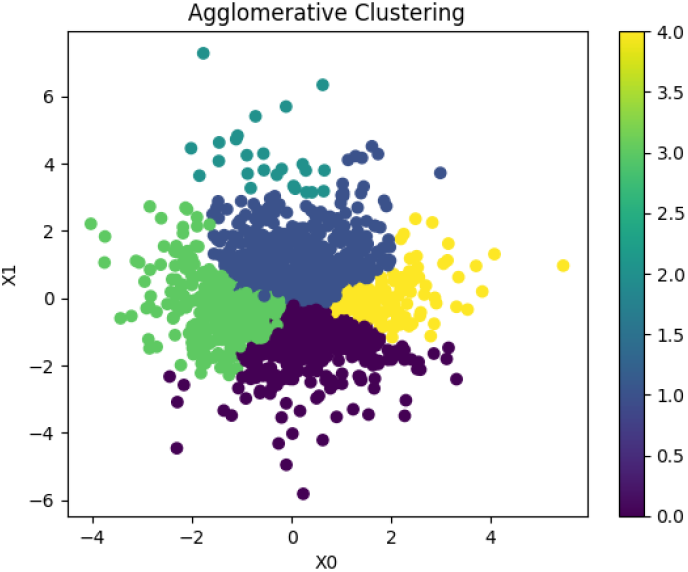
Hierarquical clustering of proteins in 25pdb based on its physicochemical properties.

On top of this, a classification strategy is implemented on which amino acids from proteins from each one of these clusters are assigned to a given CNN model built using parameters described above and a final neural network that combines the outputs from all the CNNs into a final prediction.

Following this method, amino acids weren’t distributed evenly among the CNN models. Figure 9 provides a representation of the amount of data assigned to each model. Here is important to notice that while some clusters have a high number samples while others have a relatively low number of samples. It would be interesting to add more proteins to boost training in models assigned to these low populated clusters.

**Figure 9.**
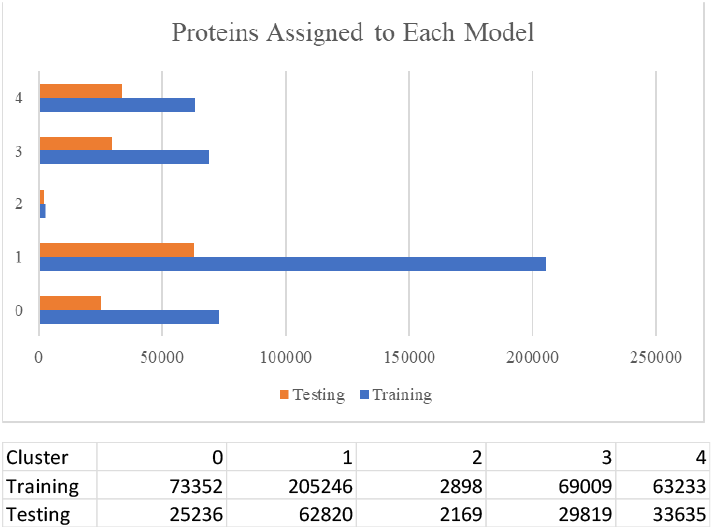
A representation of the number of amino acids assigned to each CNN model.

The overall mechanism of SSPCP can be seen in Figure 10. Protein features are extracted using the method explained above. Then a clustering strategy is implemented on top of a two component PCA. According to the cluster assigned to each protein, data from its amino acids (extracted from PSSM using the same method as described in previous strategies) is sent to a particular CNN model. Finally, a neural network will combine the results of all these independent with the assigned cluster and predict a final secondary structure.

**Figure 10.**
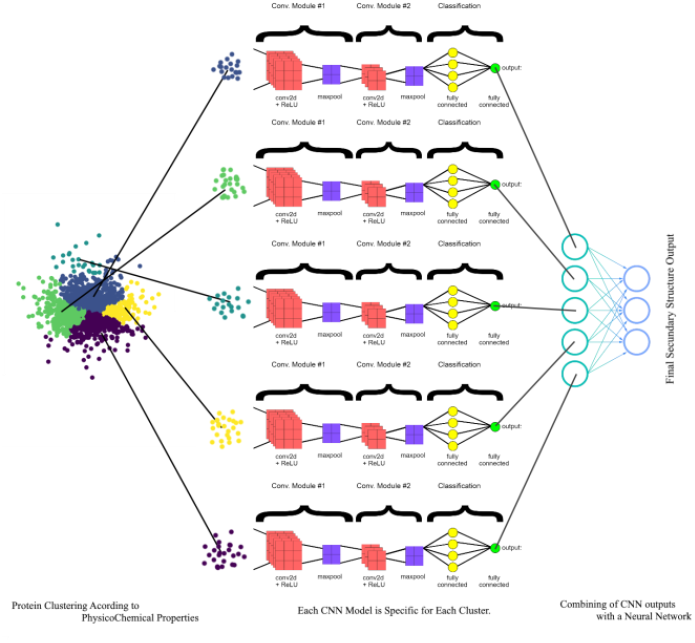
Schematics of SSPCP.

Here something interesting happens. When these highly specific CNN models are combined together, the final predicted results increase slightly in performance. The increase wasn’t amazing, but by improving training data and performing parameter fine-tuning for each of the CNN models there is expectation that these values might surpass similar methods. Also, there is possibility of using other methods concurrently with CNN models. We tried LSTM but outcome was not better.

Using a neural network, the results outperform other methods achieving **70.91%** Q3 score.

## IV. Pratical Application

For consolidation of work a practical application of SSPCP was applied to an independent protein. The protein of choice was the novel PET hydrolase from *Ideonella sakaiensis* 201-F6 which is able to hydrolyse polyethylene Terephthalate. In the Annexes the results are shown in detail. SSPCP was able to correctly predict structure for 73% of the amino acids which are highlighted in green. The main mistakes occurred when the group of amino acids change from one class to another, which seems to happen due to lack of training and possible not enough information retrieval of the proximate amino-acids. Expectedly, these results would improve with better training sets. Also, by using different classification strategies such as LSTM combined with CNN for each one of the clusters might be useful to retrieve different types of information for the correct classification of each amino acid.

## V. Discussion

For the SVM-RBF classifier the best parameters were C: 10.0 and gamma: 0.001 when applied to a subset of 100 proteins. As reported in previous works w=13 was the best choice for PSSM neighbourhood length achieving 0.75 Q3 score in the same subset with 100 proteins. During the implementation there was expectation regarding whether each cluster would be highly correlated with a given type of secondary structure, however that correlation was not found in data. All clusters had a significantly lower number of strands. Here is important to notice that maybe for future works it would be important to find datasets with provide more samples for that class, since there is a huge discrepancy between expected helix and coil classes and that can limit the performance of classifiers. It was not possible to find a direct correlation between the hierarquical clustering of features extract from PSSM matrix and secondary structure.

By using CNN, the final classification Q3 score on testing dataset was of 70.11%. By using LSTM, the final classification Q3 score on testing dataset was of 69.25%.

Ensemble CNN and LSTM provided a final classification Q3 score on testing dataset was of 70.71%. This suggests combining multiple kinds of classifiers might provide more accurate ways of predicting secondary structure, since there can be multiple reasons why these conformations are present in each amino acid.

Secondary Structure of Physicochemical Clustered Proteins (SSPCP) was proposed. It is a specific method that combines multiple CNN models trained accordingly to a previous protein feature clustering and a combining neural network. By using SSPCP, the final classification Q3 score on testing dataset was of 70.95%.

The rationale behind SSPCP is that amino acids from proteins which have similar physicochemical characteristics should have the same secondary structure prediction for similar amino acids, but amino acids from differing proteins might have different structures. In this method amino acids weren’t distributed evenly among the CNN models. It is important to notice that while some clusters have a high number of samples, others have a relatively low number of samples. It would be interesting to add more proteins to boost training in models assigned to these low populated clusters.

## VI. Future Work

For future related works it would be important to experiment with proteins which provide more coil classes, since with current dataset a very small part of proteins have this structure.

We provide an interesting rationale. However, it demands further exploration. It is important to experiment more reliable techniques for protein clustering and to explore which model might perform better for each protein subgroup. Here we implemented only CNN, but the models can be diversified inside SSPCP in order to get more accurate classifiers for each subgroup.

Also, a larger dataset might be very useful for training the SSPCP, since now each model will receive less and more accurate data. Some authors combine multiple datasets in order to have more reliable training data, that can and should also be done here.

## VII. Limitations

During the replication of previous work, some problems were found. It is not clear how to the authors converted protein structure into amino acid secondary structure for prediction. It was assumed DSSP was the method used, but it might not be. Also, it was not clear how authors sliced the dataset in order to transform 1673 proteins into 887 proteins, neither how 887 proteins provide 134853 amino acids, since protein size varies too much. On top of that, the precise structure of each model might not be exactly the same because some details were not accurate enough, for example number of neurons on classification and fully connected layers was not referred. We tried to replicate models as close as we could, but these details might help explain discrepancies between results found and the reported performances.

Also, the context of this project had limited time and thus it was not possible to deeply explore all the parameters and variables and proposed more improvements. In order to achieve better and more accurate conclusions access to more computational power would be needed. In this work Swissprot was used in order to calculate PSSM matrices from PSIBLAST, however there are other choices that might be used in the previous works such as non-redundant database, but it takes more than 400 GB of space and each calculation would take a lot more time. On top of that, note that some tests might take multiple hours to run. With more resources it would be possible to have more accurate PSSMs, find better models and train with larger datasets.

## VIII. Annexes

SSPCP Secondary structure prediction of Chain A of PET hydrolase from Ideonella sakaiensis 201-F6 PDB DOI: 10.2210/pdb5XG0/pdb

Line 1 Amino-acid Sequence Line 2 DSSP Structure from PDB Line 3 Predicted Structure

1:AMNPYARGPNPTAASLEASAGPFTVRSFTVSRPSGYGAGTVYYPTNAGGTVGAIAIVPGYTARQSSIKWWGPRLASHGFVVITIDTNSTLDQPSSRSSQQMAALRQVASLNGTSSSPIYGKVDTARMVMGWSMGGGGSLISAANNPSLKAAAPQAPWDSSTNFSSVTVPTLIFACENDSIAPVNSSALPIYDSMSRNAKQFLEINGGSHSCANSGNSNQALIGKKGVAWMKRFMDNDTRYSTFACENPNSTRVSFRTANCS

2:CCCCCCCCCCCCHHHHHCCCCCCCEEEEECCCCCCCCEEEEEEECCCCCCEEEEEEECCCCCCHHHCCCHHHHHHHHHCEEEEEECCCCCCCHHHHHHHHHHHHHHHHHHHCCCCCCCCCCEEEEEEEEEECHHHHHHHHHHHHCCCCCEECCECCCCCCCCCCCCCCCEEEEEECCCCCCCCCCCHHHHHHCCCCCCEEEEEECCCCCCCCCCCCCCHHHHHHHHHHHHHHHHCCCHHHHHHHHCCCCCCCEEEEEECCC

3:CCCCCCCCCCCCCHCHCCCCCCCCCCEEEEEECCCCCEEEEEECCCCCCCCEEEEEECCCCCCHHHHHHHHHHHHHCCEEEEEEECCCCCCCCCCCHHHCHHHHHHHHHHHHHHHHHHHHCCCCCEEEEEECHHHHHHHHHHHCCCCCCEEEEECCCCCHHHHCCCCCHEEEEECCCCCECCHHHHHHHHHHHCCCCCCEEEEECCCCCECCCCCCCHHHHHHHHHHHHHHHHHCCCCCCCEEEECCCCCCCCCCCCCCCC

Predicted C, expected E: 25

Predicted H, expected C: 23

Predicted C, expected H: 12

Predicted E, expected C: 5

Predicted E, expected H: 4

Match = 194/263 = 0.73

Mismatch = 69/263 = 0.26

## Notes

### Competing Interest Statement

The authors have declared no competing interest.

